# Strand-switching mechanism of Pif1 helicase induced by its collision with a G-quadruplex embedded in dsDNA

**DOI:** 10.1101/2022.01.11.475363

**Authors:** Jessica Valle-Orero, Martin Rieu, Phong Lan Thao Tran, Alexandra Joubert, Jean-François Allemand, Vincent Croquette, Jean-Baptiste Boulé

**Affiliations:** Laboratoire de physique de L’École Normale Supérieure de Paris, CNRS, ENS, Université PSL, Sorbonne Université, Université de Paris, 75005 Paris, France; Institut de Biologie de l’École Normale Supérieure de Paris (IBENS), École Normale Supérieure, CNRS, INSERM, Université PSL, 75005 Paris, France; Structure et Instabilité des Génomes, Museum National d’Histoire Naturelle, INSERM, CNRS, Alliance Sorbonne Université, 75005 Paris, France; ESPCI Paris, Université PSL, 75005 Paris, France

## Abstract

G-rich sequences found at multiple sites throughout all genomes may form secondary structures called G-quadruplexes (G4), which act as roadblocks for molecular motors. Among the enzymes thought to process these structures, the Pif1 DNA helicase is considered as an archetypical G4-resolvase and its absence has been linked to G4-related genomic instabilities in yeast. Here we developed a single-molecule assay to observe Pif1 opening a DNA duplex and resolving the G4 in real time. In support of former enzymological studies, we show that the helicase reduces the lifetime of G4 from hours to seconds. However, we observe that in presence of a G4, Pif1 exhibits a strong strand switching behavior, which can lead to Pif1 escaping G4 resolution, depending on the structural context surrounding the substrate. This behavior is also detected in presence of other roadblocks (LNA or RNA). We propose that the efficiency of Pif1 to remove a roadblock (G4 or other) is affected by its strand switching behavior and depends on the context surrounding the obstacle. We discuss how this switching behavior may explain several aspects of Pif1 substrate preference and affect its activity as a G4 resolvase in vivo.

## Introduction

G-quadruplexes (G4) are nucleic acids (NA) secondary structures that may form when four stretches of successive guanines appear consecutively along the primary sequence. Under appropriate ionic conditions, guanines of each stretch line up to form planes (G-quartets) with four coplanar guanines interacting via Hoogsteen hydrogen bonds. Between these planes, cations such as K^+^ help stabilize the structure, which may form intra- or intermolecularly and have different folding patterns (typically referred to as parallel, anti-parallel, or mixed structures [1, 2]). Formation of nucleic acids such as G4 are suspected to act as roadblocks for the replication machinery, causing fork pausing and potentially promoting DNA breakage [3–6]. Despite causing potential roadblocks for molecular motors, there is increasing evidence that many G4 forming sequences were selected for during evolution, suggesting a role for encoding structural information within DNA at the expense of evolving protein motors able to remove these stable structures during DNA replication or repair [7–10]. A well studied G4-resolving motor is the *Saccharomyces cerevisiae* DNA helicase Pif1, a multifunctional helicase which plays multiple roles in nuclear and mitochondrial genome stability, including resolution of G4 structures [11]. Pif1 exhibits a 5’-to-3’ polarity and translocates on ssDNA while displacing complementary DNA, RNA or proteins [12–15]. Both bulk and single molecule studies have shown that Pif1 has a high affinity for DNA but unwinds dsDNA with low processivity [14, 16, 17] and that dimerization of Pif1 is necessary for efficient translocation on DNA [18]. In addition, single molecule studies have led to the proposition of a mechanism named ”patrolling”, which consists of Pif1 reeling ssDNA while anchored at a ss-dsDNA junction [16]. In a separate study however, this activity was shown to be rare compared to translocation [19]. Interestingly, repetitive unwinding of a substrate by Pif1 has been a regular theme in Pif1 single molecule studies, but the underlying mechanism has remained unclear. Therefore, a complete description of Pif1 translocation mechanism, its substrate preference and its mechanism of G4 unwinding is still lacking.

A main experimental challenge for studying intramolecular G4 unwinding in single molecule assays is to be able to control the formation and lifetime of G4 structures. We recently developed a single molecule assay allowing embedding a G4 structure within a dsDNA molecule, mimicking a dsDNA fork, a situation which may occur in the cell during replication or at a gene promoter [20]. Using this assay, we sought to gain insights into the mechanism of G4 resolution by the *Saccharomyces cerevisie* Pif1. Here, we provide a real-time visualization of Pif1 translocating along dsDNA and interacting with a stable G4 structure taken from the promoter of the human cMYC gene. We monitored the position of the fork with few nanometers resolution, allowing us to follow the helicase at base pair resolution as it encounters the embedded G4. We show that in presence of an available single stranded DNA in the vicinity of the enzyme upon collision with the G4, Pif1 exhibits a strand switching behavior. In this situation, G4 resolution depends therefore on the ability of Pif1 to collide multiple times with the G4 after several rounds of strand switching. In absence of a ssDNA strand in the vicinity of the enzyme, the enzyme pauses in front of the obstacle until it removes the G4 and resumes translocation. Finally, we show that Pif1 exhibits strand switching in presence of other obstacles, namely LNA:DNA or RNA:DNA hybrids and obtained the probability rate of this mechanism for each substrate. Our results suggests that the ability of Pif1 to remove a G4 (or other obstacles) is affected by the availability of a single stranded DNA allowing strand switching. We discuss the *in vivo* relevance of our observations in the context of Pif1 substrate preference and *in vivo* roles.

## Materials and Methods

### DNA substrate

All oligonucleotides were bought from Eurogentec (Seraing, Belgium) or Integrated DNA Technologies (Leuven, Belgium). The tethered DNA substrate was a 310-bases long molecule comprised of a 87-bp hairpin region, a 6-nucleotide loop, and two single-stranded (ss) DNA handles allowing attachment to the surface and the magnetic bead respectively. We designed two versions of the substrate that differ in the way Pif1 encounters the G4: one where the G4 collision occurs when opening the hairpin (*LagG4)* and one where Pif1 collides with the G4 while the hairpin is closing behind the translocating helicase (*LeadG4). LagG4* contains the Cmyc-PU27 G4-motif (GGG-GAG-GGT-GGG-GAG-GGT-GGG-GAA-GG) in the center of the hairpin region, between the 5’ extremity and the loop. *LeadG4* contains the same motif, this time between the loop and the 3’ extremity. Both hairpins are formed via ligation of three oligonucleotides, Oligo5_*X*_, Oligo3_*X*_ and OliLoop, where X stands for *LeadG4* or *LagG4*. The 5’ ss-handle of the hairpins is complementary to a 58-base 3’-DBCO modified oligonucleotide (OliDBCO) attached to the surface. The 3’-end of the hairpin is complementary to a 57-base oligonucleotide (OliBiotin), which contains two biotin modifications at its 5’-end. Between the region complementary to OliDBCO and the hairpin region, seven bases (poly-dT) of single-stranded DNA allow the binding of Pif1 to the substrate. OliRNA and OliLNA are 34-base oligonucleotides that are complementary to the *LeadG4* hairpin region containing the G4 motif. OliLNA is composed of 25 bases of DNA and 9 bases of LNA on its 3’ extremity. All oligonucleotide sequences are available in Supplementary Table S1 and Table S2. A schematic of the single molecule substrate is available on Supplementary Figure S1.

### Magnetic tweezers

All our measurements were carried out using magnetic tweezers, which is a single-molecule force spectroscopy technique. They consist in measuring the extensions of single DNA hairpin molecules tethered to a surface on one end and on a magnetic bead on the other end, in function of the force applied to the magnetic bead the DNA hairpin is tethered to. The force is provided by a couple of permanent magnets and is proportional to the magnetic field gradient [21–24]. The force is calibrated using the fluctuation dissipation theorem and can be modified by changing the distance of the magnets to the surface. Our force range expands from 0 - 22 pN, with a variability from bead to bead of 10%, due to the heterogeneity of their magnetization. The magnets are positioned with a DC-motor (Physik Instrumente, M-126K070) to modulate the force. Up to 100 magnetic beads are imaged using a CMOS camera. Their 3D-position is inferred in real-time with few nanometer precision using tracking techniques.

### Bead preparation

Purified DNA hairpin was first hybridized with OliBiotin during 30 minutes by mixing both molecules at 5 nM in passivation buffer (140 mM NaCl, BSA 1%, Pluronic F-127 1%, 5 mM EDTA, 10 mM NaN_3_, pH 7.4). 5 *μ*L of streptavidin coated Dynabeads MyOne T1 (Thermofisher) were washed three times with 200 *μ*L passivation buffer. The hybridized substrate was diluted to 200 pM and 2 *μ*L of this solution was then incubated 10 minutes with the beads in a total volume of 20 *μ*L of passivation buffer. The beads were then rinsed three times with passivation buffer in order to remove unbound DNA. All the reactions were performed at room temperature.

### Pif1 helicase expression and purification

The sequence coding for the nuclear form of yeast Pif1 (amino acids 40–859, in this work referred to as Pif1) fused at its N-terminus to a 6-histidine tag was cloned into vector pET28 (Novagen) and transformed into E. coli BL21(DE3) strain (Novagen). Cells were cultivated in Luria Broth media and protein expression was induced for 16h at 18°C by addition of 0.2 mM IPTG at 0.8 OD_600_, followed by another induction with 0.2 mM IPTG for 4h at the same temperature. The cell pellet was resuspended in buffer P (20 mM sodium phosphate buffer pH 7.5, 500 mM NaCl, 1 mM TCEP and 10% glycerol) and sonicated on ice (40% amplitude, pulse 40%, 10 min, 2s on/2s off) to lyse the cells. The lysate was subjected to ultracentrifugation at 30 000g for 1h. The supernatant was applied onto a HisTrap Crude affinity column (Cytiva), washed with five bed volumes of buffer containing 20 mM imidazole and eluted with a 20 mM to 200 mM imidazole gradient. Fractions containing Pif1 were loaded onto a CHT column (Biorad) and eluted with a 20 mM to 200 mM phosphate gradient. Finally, the fractions containing Pif1 were dialized against buffer S (50 mM Hepes pH7.5, 100 mM KCl, 1 mM TCEP, 10% Glycerol), loaded onto a strong cation exchange column (MonoS, Cytiva) and eluted with a 100 mM to 1M salt gradient. Fractions were separated by electrophoresis on denaturing polyacrylamide gels. Fractions containing pure Pif1 as assessed by Coomassie staining of the gels were pooled, concentrated and stored at –80°C in 25 mM HEPES pH 7.5, 250 mM KCl, 0.5 mM TCEP, 50% glycerol.

### Single molecule Pif1 measurements

Our experiments began with the preparation of a microfluidic chamber coated with OliDBCO (Supplementary Table S2). To do so, a 40-*μ*L droplet containing 100 nM OliDBCO and 500 mM NaCl was incubated for two hours on a azide-functionalized coverslip (PolyAn, Berlin, Germany) at room temperature. The coverslips were then rinsed with passivation buffer and assembled into a 10-*μ*L microfluidic chamber. After the chamber was attached to the magnetic tweezer setup, 1 *μ*L of the bead solution was introduced into the cell, filled with passivation buffer and incubated for 10 minutes. Excess unbound beads were washed out by flowing passivation buffer into the cell. Measurements started by applying repetitive force cycles between 4 pN and 19 pN to the tethered molecules. Only molecules that showed an instantaneous unzipping at 19 pN (corresponding to an immediate increase of extension of 100 to 120 nm), followed by a complete re-zipping when the force was lowered to 4 pN, were kept. This signature allowed us to select molecules attached as expected as a result of annealing between OliDBCO and the hairpin molecule, and not through non specific interactions. Once the well-attached hairpins were identified, the microfluidic chamber was equilibrated with 200 *μ*L G4Pif1 buffer (10 mM Tris-HCL (pH 7.5), 100 mM KCl, 3 mM MgCl_2_), where potassium enhances the stability of G4 structures and magnesium is necessary for Pif1 activity [25]. Then, a 7-base oligonucleotide (Oli7) at a concentration of 10 nM, complementary to the hairpin loop was injected in the solution (Supplementary Table S2). We then applied repetitively the following force cycle: 19 pN for 8s (hairpin unzipping and Oli7 hybridization), 7 pN for 15s (hairpin rezipping blocked by Oli7), and 5s at 4 pN (expulsion of Oli7 and full rezipping). At 19 pN the hairpin fully unzips, and rezips at 7 pN. However, the hybridization of Oli7 to the loop at 19 pN temporarily prevents the closing of the hairpin, even at 7 pN. This low force regime with extended ssDNA thanks to Oli7 enables the formation of the G4 structure. This is probed, under the absence of Oli7, as a rezipping of only half of the hairpin (blockage at G4 position) when the force is lowered to 7 pN. See a detailed description of the protocol of G4 formation and detection in [20]. Once all G4 were formed and before injecting Pif1 we performed the following force cycle protocol: we increased the force to 19 pN for a full opening of the hairpin, followed by a closing at 7 pN, where again a blockage at the G4 position was observed; we then increased the force to 11 pN for a few seconds to determine the extension of the partial closed hairpin and thus the position of the G4, lowered the force to 4 pN in order to close the hairpin around the G4 (embedded G4), and finally we increased the force to 11 pN (Figure 1B-red). From this point on in the experiment, the force was kept at 11 pN and Pif1 was added to the microfluidic chamber at a concentration of 6 nM and 1 mM ATP Lithium salt (Roche) in G4-Pif1 buffer. A flow of 5 *μ*L/min of this Pif1/ATP was kept constant throughout the experiments. In the case of the experiments with RNA:DNA and LNA:DNA, addition of Oli7 was omitted so that G4 was not induced. The same force protocol was applied as above until the hybridization of OliRNA and OliLNA took place, which was observed as the rezipping of only half of the hairpin at 7 pN (same position as the G4).

**Figure 1.**
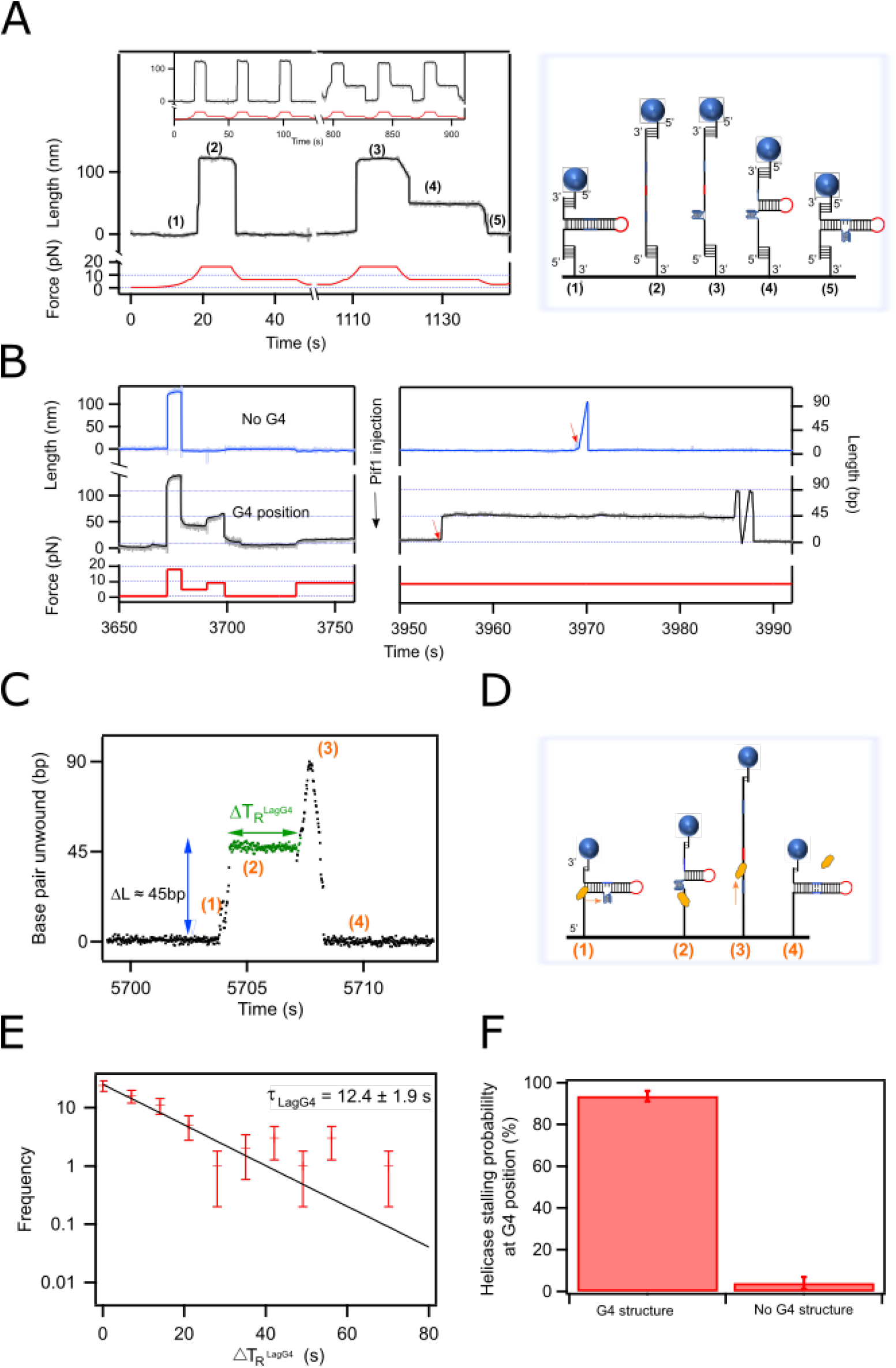
G4 formation on a *LagG4* hairpin assay. **A) G4 prevents the re-zipping of the hairpin.** Force-extension curves and sketches of our hairpin assay. Cycles of 19 - 7 pN show the unzipping (2) and re-zipping (1) of the hairpin respectively. However, when G4 is formed, a partial re-zipping is observed at 7 pN (4); lowering the force even more (4 pN), forces the hairpin to close while encircling the G4 structure (5). **B) Pif1 binds to *LagG4* hairpin.** Red trace: Force protocol to test G4 presence and interaction with Pif1. The force is kept constant at 11 pN when Pif1 is injected and throughout the measurement. Blue trace: G4 has not formed, as observed by the lack of blockage when the force is lowered to 7 pN. Pif1 binds to the DNA (red arrow) and translocates throughout the hairpin without being stalled by the G4-structure. Black trace: G4 has formed, as observed by a blockage at 7 pN. Pif1 attaches to the hairpin (red arrow) and unwinds it until is stalled at the G4 position **C) Pif1 is stalled by the G4.** Pif1 opens about 30 base pairs and translocates within the G4 motif about 15 additional nucleotides (accounting for 45 unwound bp), until it gets stalled by the G4 for a period of time 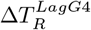 (green trace, steps 1 to 2), time needed for Pif1 to resolve G4. As Pif1 continues travelling on the opposite strand (step 3), the hairpin closes back behind the helicase (step 4). **D) Sketch of *LagG4* hairpin** Provides a cartoon simulating the extension of the hairpin throughout the different steps from C as Pif1 travels through it (yellow). **E) Distribution of resolving time of G4 quadruplexes by Pif1**. The resolving time shows a single exponential characterized by a time constant *τ*_*LagG*4_ = 12.4 ± 1.9 s. **E) Statistics.** Left Bar: Probability Pif1 is stalled by G4 after it had folded. Right Bar: Probability there is a blockage at the G4 position when G4 was not present in the previous cycles.

### Data collection and analysis

Molecule extensions and forces were collected in real-time using Xvin, an interactive data visualization software developed in-house in C/C++ [26], at an acquisition frequency of 50 Hz and at a constant temperature of 25°C. Extension graphs were generated using IgorPro (Wavemetrics, Lake Oswego, OR, USA). To interpret the extension of the hairpins as positions of the helicase along the sequence, extensions in nanometers were converted to basepairs. To do so, the average extension change of the hairpins upon unwinding is measured at 11 pN as the distance between the lowest point and the highest point of a full unwinding event of the hairpin by Pif1. This distance is divided by 90 bp (length of the hairpin + loop), which gives the length corresponding to the unwinding of one base pair (see Supplementary Methods). All traces show both the raw data of extension (in nm or bp) vs time (in seconds) (light points) and the filtered data (dark curve). The latter was obtained using a non-linear filter slope algorithm [27].

Individual stalling events during hairpin unwinding and rezipping were manually detected using Xvin. We considered that there was a pause in either unwinding or rezipping when the position of the bead stayed within a window of 5 nm during a time larger than 0.3 s. In comparison, the bead moved at a velocity of ≃ 100 nm/s upon enzyme translocation, and thus would cross a 10-nm window in 0.1 s. Empirically, all stalling events occurred close to the G4 position in the sequence (50 ± 5 bases) (see Supplementary Methods and Figure S13). Stalling times Δ*t′* correspond to the times elapsed between the start and the end of individual stalling events. The resolving times 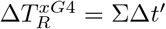 (with x being *Lead* or *Lag* substrates) correspond to the total time spent in stalling events by the enzyme before it bypasses the G4 position. Between 50 and 100 resolving times were measured for both of the substrates, and histograms were computed with a binsize of around 7s. The histograms and corresponding errors (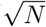, N being the counts in each bin) were fitted to single exponential distributions using IgorPro. The characteristic time rates *τ*_LagG4_ and *τ*_LeadG4_ correspond to the parameters inferred from this fitting procedure. Error bars are also computed with IgorPro using the covariance of the fit. In our studies of Pif1 with *LNA:DNA* and *RNA:DNA* hereroduplexes, we computed the resolving times as the average of the contact times of Pif1 with the heteroduplexes before the removal of the oligonucleotide, defined as 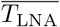 and 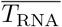 for LNA and RNA respectively. Likewise, we also obtained Δ*t*. The histograms of Δ*t* were computed with a binsize of 0.2 or 0.3 s, and fitted to single exponential distributions, from which a time constant 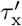 (where x is G4, LNA or RNA) was obtained.

Translocation velocities of Pif1 were measured separately during unzipping (*v*_unz_) and rezipping (*v*_z_) for all our substrates, by fitting the position of the bead with a linear function. More than 50 slopes from individual events obtained this way were then averaged to obtain the mean translocation velocities. Average velocities were converted from nm/s to bp/s as explained above and in Supplementary Methods. All these statistics are summarized in Table 1.

**Table 1.**
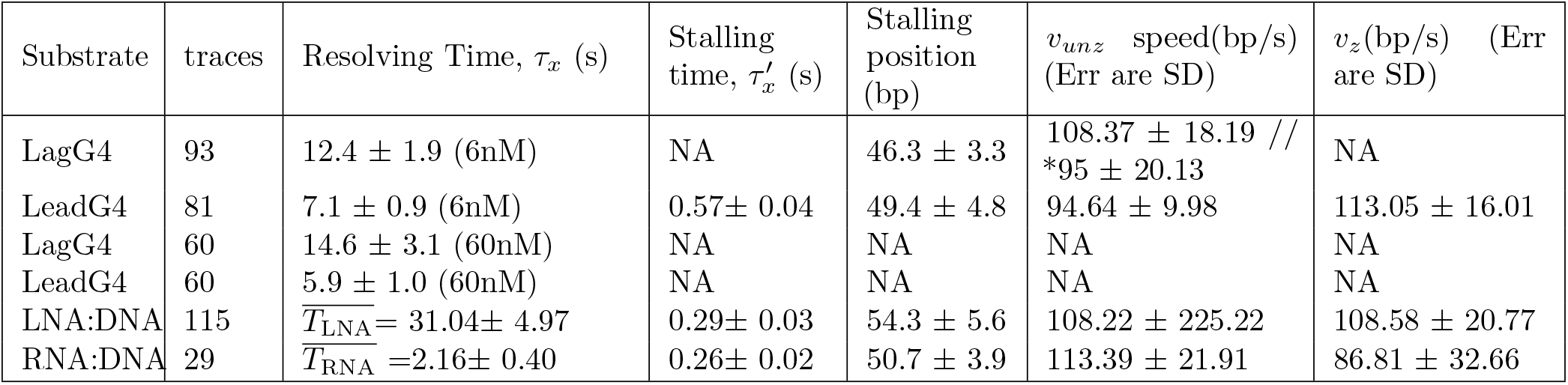
Summary of Statistics. This is a summary table of the statistics compute for Pif1 helicase during its interaction with different substrates: G4 formed in the lagging strand (*LagG4)*, or in the leading strand (*LeadG4)* at two different helicase concentration: 6 and 60 nM. As well as the dynamics of Pif1 when interacts with a hybrid complex: LNA:DNA or RNA:DNA, at the G4 position in the leading strand. In the *LagG4* susbtrate we obtained two unzipping speeds of Pif1, before interacts with G4 (first value) and after the stalling (marked with *). All speeds were measured in nm/s and converted to bp/s as described in the Supplementary materials, together with table SI1.

| Substrate | traces | Resolving Time, $\tau_x$ (s) | Stalling time, $\tau'_x$ (s) | Stalling position (bp) | $v_{unz}$ speed(bp/s) (Err are SD) | $v_z$(bp/s) (Err are SD) |
| --- | --- | --- | --- | --- | --- | --- |
| LagG4 | 93 | $12.4 \pm 1.9$ (6nM) | NA | $46.3 \pm 3.3$ | $108.37 \pm 18.19$ // *$95 \pm 20.13$ | NA |
| LeadG4 | 81 | $7.1 \pm 0.9$ (6nM) | $0.57 \pm 0.04$ | $49.4 \pm 4.8$ | $94.64 \pm 9.98$ | $113.05 \pm 16.01$ |
| LagG4 | 60 | $14.6 \pm 3.1$ (60nM) | NA | NA | NA | NA |
| LeadG4 | 60 | $5.9 \pm 1.0$ (60nM) | NA | NA | NA | NA |
| LNA:DNA | 115 | $\overline{T}_{LNA} = 31.04 \pm 4.97$ | $0.29 \pm 0.03$ | $54.3 \pm 5.6$ | $108.22 \pm 225.22$ | $108.58 \pm 20.77$ |
| RNA:DNA | 29 | $\overline{T}_{RNA} = 2.16 \pm 0.40$ | $0.26 \pm 0.02$ | $50.7 \pm 3.9$ | $113.39 \pm 21.91$ | $86.81 \pm 32.66$ |

## Results

### Single-molecule observation of Pif1 unwinding a G4-containing hairpin unveils how it resolves the secondary structure

We investigated the interaction between Pif1 and a G-quadruplex structure while the helicase unwinds dsDNA. For this purpose, we designed a hairpin of 87 base pairs which sequence contains the G4 motif of the c-Myc promoter sequence (c-Myc Pu27) [28] (see Supplementary Figure S1).

#### Formation and detection of a G4 structure embedded in a dsDNA

We formed the G4 structure in the lagging strand of a hairpin (*LagG4*) by applying repetitive force cycles in the presence of 100 mM potassium and of a 7 bp oligonucleotide complementary to the hairpin apex, as described in [20]. Figure 1A shows the changes of extension of the hairpin upon changes in the pulling force. Pulling on a closed hairpin (1) at 19 pN results in its full unzipping and consequent increase of the extension by about 120 nm (∼ 90bp) (2). When the force is lowered to 7 pN, the hairpin rezips (back to (1)). However, when G4 forms, the opened hairpin (3) only partially closes back (4) when the force is reduced from 19 pN to 7 pN, and a blockage is observed at an extension of about 50 bp, position at which the G4 motif is expected. Once formed, without Pif1, the G4 structure does not unfold spontaneously during several hours (Supplementary Figure S2). The force is then lowered to 4 pN in order to drive the full closure of the hairpin while encircling the G4 structure (5). A cartoon of our hairpin assay at the different extensions upon changes in the pulling force is shown in Figure 1A-right.

The dynamics of Pif1 unwinding was measured by tracking the position of the bead as described in [17]. For this purpose, we developed a force protocol (Figure 1B-bottom panel) that allows visualizing different extensions of the hairpin in the presence of the G4 structure, as well as the interaction of the helicase with the structure. Once a G4 was folded, the force was kept at 11 pN and Pif1 was added to the microfluidic chamber at a concentration of 6 nM in presence of 1 mM ATP. This force was chosen for two reasons: it is low enough to ensure that the hairpin remains closed, and it is high enough to increase the processivity of Pif1 larger than the size of the hairpin [17].

#### Pif1 resolves G4 structure but stalls transiently

In the absence of preformed G4 structure, as observed by the total rezipping at 7 pN, (Figure 1B-blue, left panel), Pif1 binds to the single-stranded region of the closed hairpin (red arrow) and translocates along the lagging strand in the 5’-to-3’ direction. This results in the opening of the hairpin, which is detected by the extension of the molecule, and its subsequent rezipping once Pif1 has translocated past the hairpin loop and continues to translocate along the leading strand, which is seen by a decrease of extension at the same rate. Alternatively, we also observed the dissociation of the helicase from the ssDNA, inferred from the spontaneous closing of the hairpin (Figure 1B-blue, right panel).

In the presence of a preformed G4 structure on our first construct *LagG4* (Figure 1B-black, left panel), Pif1 binds to the hairpin (red arrow) and encounters the G4 structure during hairpin opening. The unwinding traces display a characteristic pause at the G4 position (Figure 1B-black, right panel at 3955 s and Figure 1C at 5704 s, see also Supplementary Figure S3). After this pause, Pif1 resumes translocation all the way through the loop, and along the leading strand resulting in the closure of the hairpin behind the helicase. Furthermore, the absence of blockage during the rezipping of the hairpin confirms that the G4 structure was unfolded and did not reform after Pif1 translocated past it. Indeed, the absence of blockage is a signature of G4 unfolding: a force of 11 pN is too high to allow encirclement of the hairpin around a formed G4 structure [20]; if the G4 structure was still present, the hairpin would not close and we would observe a blockage at the same position as prior to Pif1 action. Another evidence for G4 unfolding is that subsequent hairpin opening by Pif1 shows no pause at the G4 position, as observed in the first passage (Supplementary Figure S5).

Figure 1C and D illustrates Pif1 unwinding the G4-containing hairpin substrate *LagG4* : first, the helicase attaches to the hairpin and translocates about 45 bases (1) before stalling at the level of the G4 (stalling position) for a varying period of time 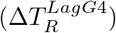 (2). After resolving the G4, Pif1 resumes translocation all the way the loop (3) and onto the other strand resulting in the full closing of the hairpin and dissociation of the helicase (4). Pif1 pausing times follow a single-exponential distribution (Figure 1E), with parameter *τ*_LagG4_ = 12.4 ± 1.9*s. τ*_LagG4_ is much smaller than the spontaneous unfolding rate of the G4 at 11 pN in the absence of Pif1 (*τ*_spontaneous_ = 27400 ± 6300 s, see Supplementary Figure S2), which indicates that Pif1 catalyzes the unfolding of the G4 structure. Thus, we referred to the time spent by Pif1 at the G4 position as the G4 resolving time 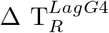. Figure 1F shows that while stalling (i.e. blockage at G4 position) is observed during almost all unwinding events in the presence of G4 structure (94%), it is almost never observed in the absence of G4 structure (4% of unwinding events). Residual events can be attributed either to detection errors or to rare events of spontaneous folding/unfolding of G4 during the time elapsed between the detection cycle (Figure 1B-left) and the unwinding events (Figure 1B-right). Overall, our results confirm that Pif1 stalling is characteristic of the presence of a folded G4 structure. Our setup thus allows us to characterize the coupled dynamics of translocation and G4-interaction of Pif1. The observation of a pause in the presence of a G4 confirms previous FRET experiments [29].

### Pif1 switches strands an resumes translocation when stalled by a G4 in a geometry where the opposite strand is available

#### Another hairpin substrate geometry allows Pif1 to interact with the opposite strand

In the geometry of the previous assay (*LagG4)*, the application of a force prevents the interaction of Pif1 with the opposite strand when it is stalled by the G4-structure (Figure 1D-(2)). Indeed, the G4 structure lies between the enzyme and the fork. Therefore, we designed a different hairpin substrate, referred to as *LeadG4*, in which the same G4 motif is located past the loop for a motor translocating in the 5’-to-3’ direction (Figure 2B and Supplementary Table S1). In this new configuration, Pif1 encounters the G4 structure after it translocated through the loop, and is located at the fork when it collides with the G4. Thus, we were able to study its interaction with a G4 structure in the vicinity of the opposite strand.

**Figure 2.**
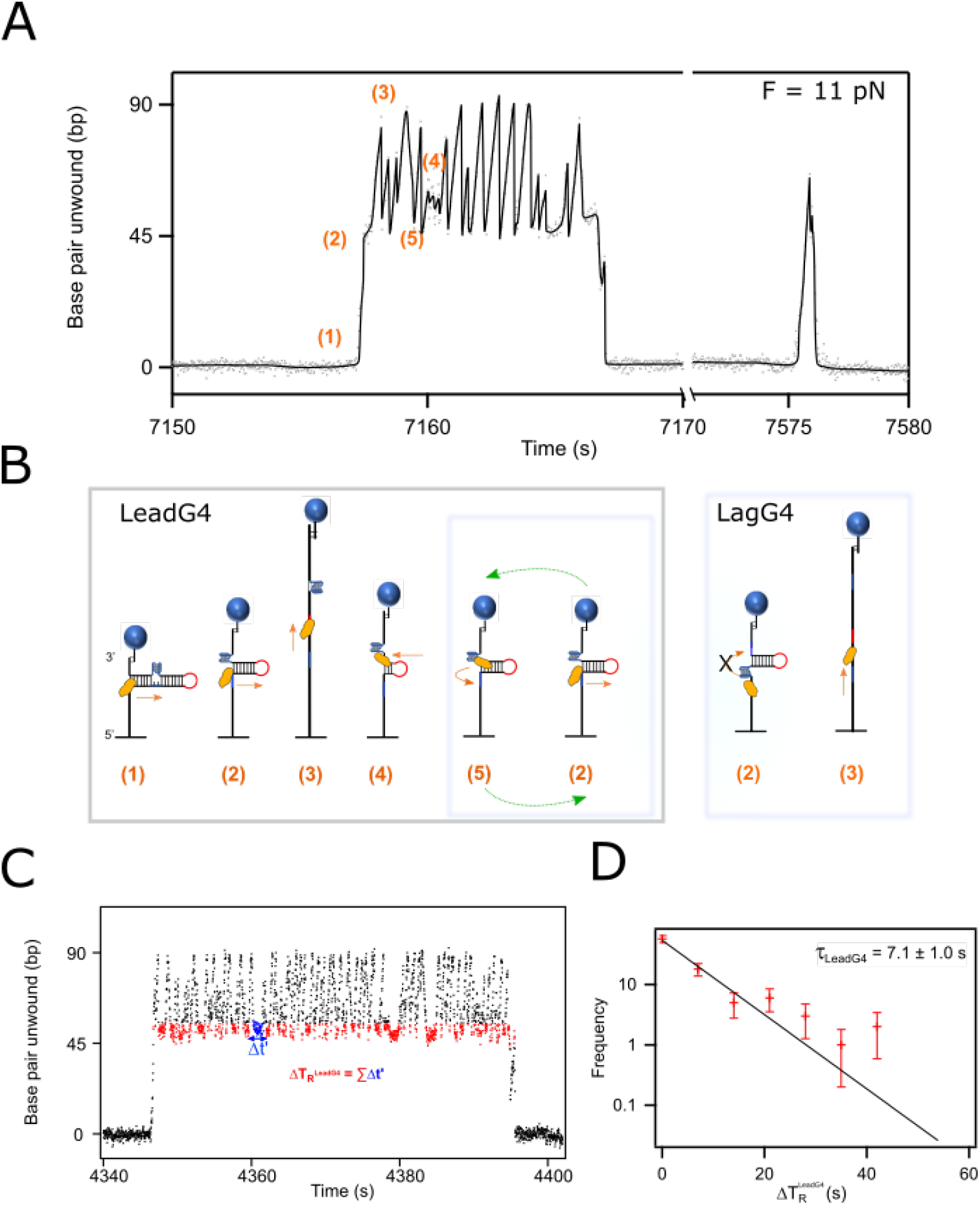
**A) Pif1 activity on LeadG4 hairpin.** Representative trace showing the collision of Pif1 when encountering the G4 structure while translocating through the hairpin. Various extensions of the hairpin are observed as Pif1 travels through it. Most relevant positions are indicated as orange numbers and described in the schematics in (B). Secondary opening of the hairpin by Pif1 after G4 resolution shows no blockage (at 7575s) **B)Sketch of *LeadG4* hairpin** Left-In the *LeadG4* assay, the helicase begins strand switching at the G4 position, visiting 5 and 2 repeatedly;right-In the *LagG4* assay, strand switching is not favored.**C) Pif1 is stalled by G4** The time Pif1 spends in contact with G4 before strands-switching, i.e. stalling time, is defined as Δ t’; the sum of these times (blue) gives the resolving time 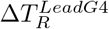, the total time Pif1 takes to resolve the G4 structure.**D) Distribution of resolving time of G4 in the *LeadG4* configuration at 6 nM of Pif1** The resolving time shows a single exponential characterized by a time constant *τ*_*LeadG*4_ = 7.1 ± 0.9 s.

To form the G4, we used the same experimental protocol as the one used for *LagG4* assay. Figure 2A shows a representative trace of the dynamics of Pif1 during its interaction with the *LeadG4* substrate containing a G4 embedded within the hairpin, at the constant force of 11 pN. In this geometry, as expected, Pif1 translocates through the hairpin past the loop (Figure 2A and B-(steps (1) to (3)), onto the other strand causing the rezipping of the hairpin until it meets the G4-structure, (steps (4) and (5)).

#### G4 structure induces Pif1 strand switching

If Pif1 behaved in the same way in this new context, we would expect a single long pause at the G4 position during the rezipping of the hairpin. Then, Pif1 would resume translocation and the hairpin would close completely. Interestingly, while we indeed observed a pause at the position of the G4 in the leading strand (stalling position, step (5)), in most cases it was followed by an extension of the molecule, i.e. Pif1 resuming translocation back towards the loop (Figure 2A). As Pif1 translocation is directional, we interpreted this different behavior as Pif1 switching strand, allowed by the proximity of the complementary strand. Then, Pif1 resumes its translocation and passes the loop until it gets blocked again at the G4 position. This induces back-and-forth movements: the hairpin loop of our substrate confines the displacement of the helicase, so that after switching direction at the G4 position, it finds itself again at the same position a few milliseconds later (Figure 2B-((5) to (2), green arrow)). After these repetitive unwinding and rewinding of the hairpin, Pif1 finally goes through the G4 location and continues its translocation resulting in the full closing of the hairpin (1). We defined this behavior as a strand-switching mechanism: Pif1 being stalled by the G4-structure can switch to the opposite strand with some probability and resume translocation in the other direction. When Pif1 finally goes through the G4 position, we see again, as in the case of *LagG4*, that the G4-structure has been resolved, since the hairpin can fully close and does not exhibit the characteristic blockage at the G4 position. Furthermore, subsequent events generally do not show any more blockages (Figure 2B and Supplementary Figures S4 and S5.

We quantified the resolving time of the G4 by Pif1 with the *LeadG4* substrate, 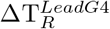, as the total time spent by Pif1 in contact with the G4 before it resolves the secondary structure (sum of the individual blue times shown in Figure 2C). Here again, 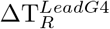 follows a single-exponential distribution characterized by a time constant *τ*_LeadG4_ = 7.1 ± 1.0*s* (Figure 2D). While it is hard to interpret why *τ*_LeadG4_ is twice smaller than *τ*_LagG4_, this decrease of *τ* is not surprising, given the difference of geometry. Indeed, in this case, the fork rezips behind Pif1. This might favor its translocation ahead compared to the other geometry, where Pif1 is unwinding the hairpin during G4 resolution. Individual stalling times (Δ*t′*, in blue in Figure 2C), also follow single-exponential distributions (Figure 3C-black) of parameter 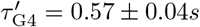. This time is representative of the probability to switch strands.

**Figure 3.**
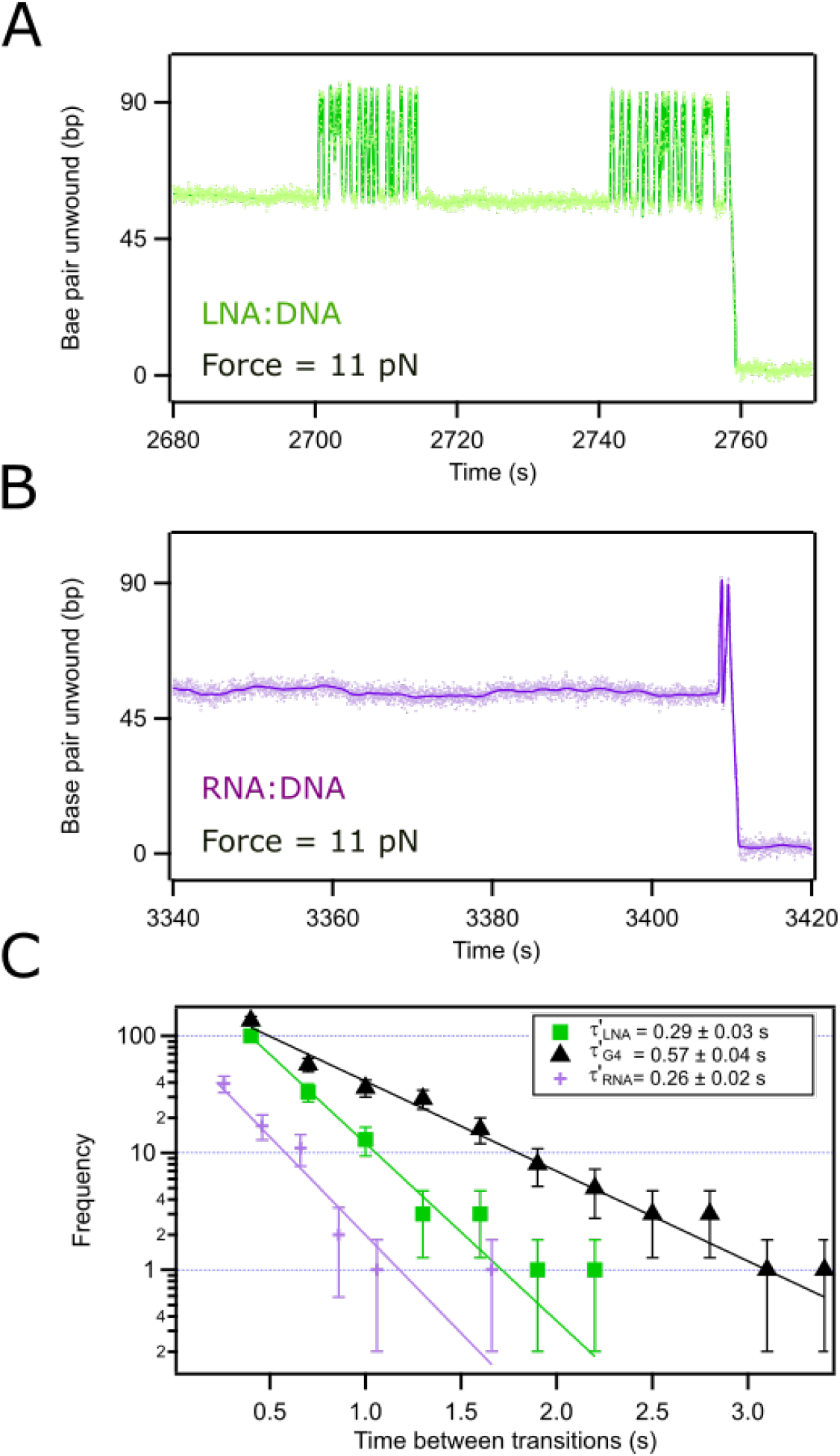
Pif1 translocates through DNA:LNA and DNA:RNA heteroduplexes. **A) Strand switching of Pif1 stimulated by LNA blockage.** Representative trace of hairpin extension in presence of Pif1 and a LNA-DNA heteroduplex. **B) Strand switching of Pif1 stimulated by RNA blockage.** Representative trace of hairpin extension in presence of Pif1 and a RNA-DNA heteroduplex.**C) The stalling time** Δ**t’ for different obstacles: G4, LNA, or RNA.** The data here corresponds to the frequency of events in log scale *vs* time-between-transitions Δt’. The probability rate obeys a single exponential distribution with a time constant 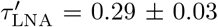 (green), 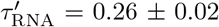 (purple), and 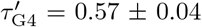 (black) for LNA, RNA and G4 respectively. In the case of G4, Δt’ includes all times measured at 6 and 60 nM, since differences were not significant.

### Pif1 exhibits strand switching with obstacles other than G4

#### Visualization Pif1 colliding with hetero-duplexes

To check whether the strand-switching mechanism observed in the previous section is induced by a specific interaction with a G4, we replaced the G4 obstacle by RNA:DNA and LNA:DNA heteroduplexes localized at the same position in the sequence as the G4 motif in the leading strand configuration (Supplementary Figure S7 and S8). Here, to avoid the formation of the G4 structure the 7 bp oligonucleotide (Oli7, see Material and Methods) was not injected into the microfluidic chamber during the experiment. Instead, we introduced 100 nM of a 34 bp LNA or RNA oligonucleotide complementary to the G4 sequence motif (Supplementary Table S1) and applied opening and closing cycles. The hybridization of the oligonucleotide to the hairpin can be detected by the blockage of the hairpin at the same position as the blockage induced by the G4 (Figure 3A and B).

#### LNA-DNA and RNA-DNA Heteroduplexes also induce Pif1 strand-switching

Once the LNA or RNA oligonucleotides hybridized, we monitor Pif1 translocation as described above. As shown in Figure 3A,B Pif1 translocation traces also display a strand-switching behaviour when Pif1 reaches the heteroduplexes. After several back and forth events, Pif1 removes the LNA or RNA oligonucleotides and resumes translocation, as indicated by the full rezipping of the hairpin.

In the case of the LNA:DNA heteroduplex, some pauses in the back-and-forth mechanisms are observed before the removal of the oligonucleotide (Figure 3A). We interpreted these pauses as the waiting time between the unbinding of the helicase from the hairpin that did not manage to remove the obstacle and the binding of another helicase. Indeed, these pauses become shorter when the enzyme concentration in solution is increased by a factor 10 (see also Supplementary Figure S6). In the case of RNA:DNA hybrid (Figure 3B), Pif1 manages to remove the RNA oligonucleotide 94% of the times before unbinding. This is not surprising considering that Pif1 processivity is enhanced on RNA:DNA substrate, and hence it preferentially translocates through the DNA strand while displacing the RNA strand [14, 16]. The resolution times were also computed as the average of the contact times of Pif1 with the heteroduplexes before their removal. We found 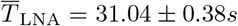 and 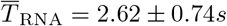. The high resolving rate 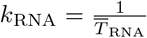 compared to the one for G4 and LNA corroborates previous results pointing the efficiency of Pif1 for unwinding DNA:RNA hybrids.

Thanks to the spatial resolution of the experimental setup it was possible to determine that strand switching and stalling occur at the start of the heteroduplex and not on further positions along the hairpin. The stalling position for all our substrates was detected at around 50 opened bases, with a resolution of 5 bases (see Table 1). This indicates that the limiting resolving step is the opening of the first bases of the heteroduplexes. Once the heteroduplexes are partially open, Pif1 proceeds smoothly with their unwinding.

The probability to switch strand was quantified by the stalling times at the obstacle position (pause time between rezipping-unzipping events) as Δ*t′* as shown in Figure 3C (also Figure 2C-blue). These times follow a single-exponential distributions of parameters 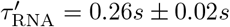 and 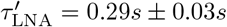 for RNA and LNA respectively, twice smaller than for G4 given as 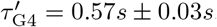.

Overall these results indicate that the strand-switching mechanism of Pif1 is not G4 specific. When Pif1 translocates along DNA and run onto a G4 or an heteroduplex, the resolution of the obstacles is dictated by a competition between three dynamic processes: the resolution of the obstacle, the escape through strand-switching, and the unbinding of the enzyme.

## Discussion

In this work, we investigated the behavior of the *Saccharomyces cerevisiae* Pif1 helicase when it unwinds a hairpin containing a G4 structure. We used the G4-detection protocols developed in our previous study [20] to form and detect the G4 within the hairpin context, and thus to compare the behavior of Pif1 in its presence and its absence. The setup allows us to visualize the translocation of the helicase along the dsDNA hairpin, its transient stalling at the position of the G4 in the sequence, and the resolution of the structure. Our results show that Pif1 considerably shortens the G4 lifetime, from hours to seconds (Supplementary Figure S2). This confirms previous results showing that Pif1 actively promotes the unfolding of G4 structures and does not wait for the G4 to spontaneously unfold in order to resume translocation [29, 30].

Two different substrates were used in this study, one in which the G4-motif is located on the translocation strand of Pif1 during unzipping (*LagG4)*, and one in which the G4-motif is located on the translocation strand of Pif1 during the rezipping of the hairpin (*LeadG4)*. For both substrates, the behavior of Pif1 displays pauses when it meets the G4, described by a stalling time Δ*t′*, that follow single exponential distributions. When the opposite strand is not available to the enzyme (*LagG4)*, the traces display a unique stalling of typical time *τ*_*LagG*4_ = 12 s, followed by the resolution of the G4 secondary structure. When the opposite strand is in close proximity of the enzyme (*LeadG4)*, we observe a short stalling time of 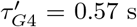 that is not sufficient to resolve the G4, but reveals a strand-switching mechanism. However, in our hairpin configuration, the strand switching promotes further interactions with the G4 and the sum of all stalling times 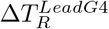 (during which Pif1 is close to the G4) before the resolution of G4 also follows an exponential distribution of typical time *τ*_*LeadG*4_ = 7 s.

Our results indicate that Pif1 can undergo two competitive mechanisms when it stalls at the G4 position. Either it switches strands and escapes the obstacle, with rate 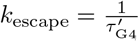, or it catalyzes the G4 resolution, and resumes translocation in the original direction, with rate 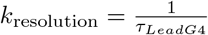. In our case, *k*_escape_ is 12 times larger than *k*_resolution_ and Pif1 switches strand most of the time. Due to the presence of the loop in the substrates, which confines the geometry of the DNA substrate, Pif1 stalls again at the G4 position a few seconds later, and so on until it unfolds the G4.

In previous single-molecule FRET experiments with a substrate mimicking a replication fork with a G-quadruplex on a free 5’ single-stranded flap [31], Zhang *et al.* observed a pause at intermediate FRET-levels between the binding of Pif1 to the fork and the subsequent unwinding of the duplex. They showed that the duration of this pause, which they called the “waiting time”, was inversely proportional to the concentration of Pif1 and thus suggested a mechanism where Pif1 is in a monomeric state when it solves the G-quadruplex and then must wait to be dimerized to perform the unwinding of the duplex. This model was supported by bulk experiments showing that Pif1 was able to dimerize on DNA substrates short as 5 bases [18]. In our case, however, the resolving time did not significantly change when we increased the injected enzyme concentration from 6 nM to 60 nM (Supplementary Figure S6 and S9, and Table 1). Although our experiments do not allow us to control the local concentration of enzyme close to the DNA-coated surface nor to test the dimerization state of Pif1, they suggest that our resolving time is not equivalent to the waiting time observed in their studies. On the other hand, it is noteworthy that we observe the same translocation velocity of Pif1 before and after the collision and the resolution of the G4 (see Table 1), which points towards Pif1 being in the same oligomeric state, whether a monomer or a dimer. Finally, the dimer/monomer transition between translocating/G4 resolving Pif1 seems to fail to account for the strand-switching behavior. If the pause between two translocation events corresponded to the time needed to bind to a second Pif1 partner, it is difficult to understand why it would be almost two orders of magnitudes lower in the case of the *LeadG4* substrate (0.57 s) with respect to the *LagG4* substrate (12 s). In our model, we understand the successive short pause times of Pif1 on the *LeadG4* substrate as the result of the competition between resolving and escaping the G4, an interpretation that is supported by the fact that the resolving time for *LeadG4* (7 s) inferred in this framework is of the same order of magnitude as for the *LagG4* substrate. Overall, our results raise questions over the current model of dynamic dimerization of Pif1.

Zhou et al., then Lu *et al.*, [16, 19], showed that the helicase could also display a patrolling behavior, where it translocates along the ssDNA while staying bound at its starting point, thus creating a ssDNA loop. In our experiments, differentiating between patrolling and standard translocation is possible by measuring the extension changes upon unzipping and rezipping. Patrolling would result in twice shorter extension changes (Supplementary Figure S11) than the one expected from the conversion between hybridized bases to ssDNA (∼ 1 nm/bp [32]). The distribution of extension changes during unzipping of the hairpin shows that no patrolling occurs in our experiments (Supplementary Figure S12). One possible cause is the small size of the ssDNA loading pad (7 bases in our experiment). Indeed, Lu *et al.* suggested that patrolling is due to a “rare oligomeric form” of Pif1, which might need a longer pad to be able to bind to its substrate.

Overall, this single-molecule assay sheds new light on the behavior of Pif1. While previous work has showed the ability of Pif1 to remove roadblocks [12, 15, 33, 34], the competitive strand-switching process that is unveiled here allows refining its role. Indeed, in the case of Cmyc-Pu27 studied here, *k*_escape_ is 12 times larger than *k*_resolution_. This means that Pif1 has 12 times more chances to switch strand than to resolve the G4. Therefore, if the substrate was a long dsDNA substrate without a confining loop as in ours Pif1 would escape the obstacle formed by G4-Cmyc-Pu27 with a probability of 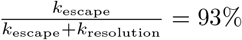. As a consequence, this strongly alleviates statements regarding the propensity of Pif1 to remove G4 structures from its translocation path, unless it is confined and forced to repetitively enter in contact with the G4, for example by a polymerase or other larger molecular complexes, as sketched in Figure 4. Strand switching would therefore appear as a regulating mechanism that prevents free Pif1 molecules to remove isolated secondary structures in the genome while activating this Pif1 feature when associated with proteins from the replisome, or as described in [33]. This mechanism would allow preserving the overall regulatory roles of G-quadruplex in the genome while preventing the stalling of replication forks. Additionally, when acting on ssDNA overhangs, absence of an available free ssDNA for strand switching would allow efficient removal of any obstacle, as was originally proposed for Pif1 action at telomeres [12]. Finally, Pif1 strand switching does not seem to be G4 specific since we observed it also during the removal of RNA:DNA and LNA:DNA heteroduplex on the translocation path of Pif1. In the case of RNA:DNA, our results confirm that the helicase removes this obstacle very easily. Interestingly, since Pif1 is not able to bind on a RNA strand [13], it may explain why the helicase exhibits higher processivity on RNA:DNA hybrids compared to dsDNA. In the case of dsDNA, the limited effective processivity of the enzyme would simply result from the competition between directional translocation and strand switching.

**Figure 4.**
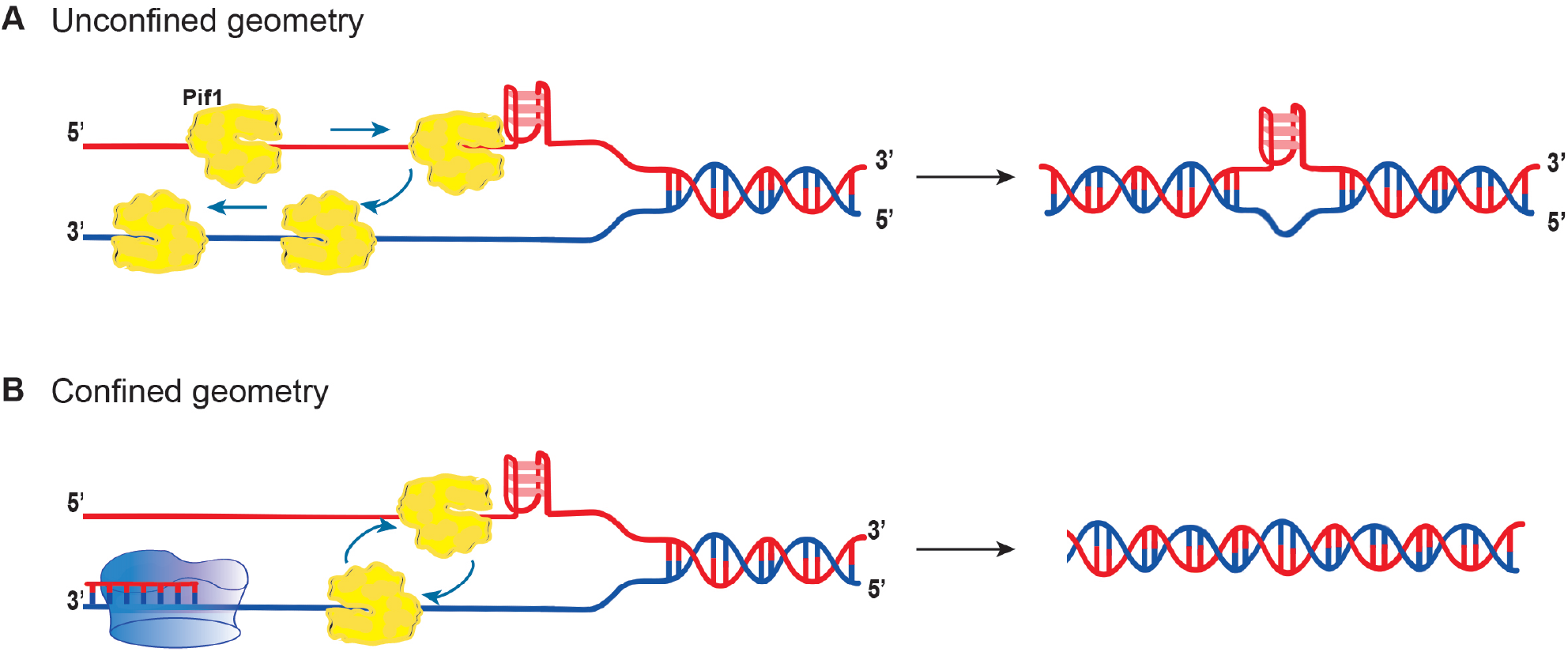
Does strand switching play a biological role? **A) Unconfined geometry.** Pif1 strand switches and is able to escape through the other strand, preventing multiple collision with the G4 structure. **B) Confined geometry.** Pif1 is trapped within a fork and cycles around by strand switching causing the reiterate collision with the G4 structure.

## Acknowledgments

Author contributions: J.V., M.R., P.L.T.T., V.C. and J.B.B. conceived the research. J.V. performed the experiments and analysis. A.J. purified the ScPif1 helicase. P.L.T.T and M.R designed and constructed the hairpins and oligos. J.V, M.R. and J.B.B participated to the writing, and all authors contributed to the editing of the manuscript. V.C. and J.B.B. supervised the research.

