## Supplementary Information for "Strand-switching mechanism of Pif1 helicase induced by its collision with a G-quadruplex embedded in dsDNA"

### SUPPLEMENTARY TABLES

| | |
|---|---|
| **LagG4** | 5'GTCTTCTTCTGTCTAATCCTTCACCGTGTCTTTTGGTCTT TCTGGTGCTCTTCGAATTTTTTTTGCAACTGTCACGATTGA CATAGCATGATAAGGGGAGGGTGGGGAGGGTGGGGAA GGATCGTACGTACAGCATCGCTTGCACTGAGAGCGCGGC CTCTCAGTGCAAGCGATGCTGTACGTACGATCCTTCCCCA CCCTCCCCACCCTCCCCTTATCATGCTATGTCAATCGTGAC AGTTGCTTTTTTTTACCGGCGCTATTAGCTTCCATACCAGCT GGCAACATCCATCATGATCCGCTACTCCCA-3' |
| **LeadG4** | 5'GTCTTCTTCTGTCTAATCCTTCACCGTGTCTTTTGGTCTT TCTGGTGCTCTTCGAATTTTTTTTGCAACTGTCACGATTGA CATAGCATGATACCTTCCCCACCCTCCCCACCCTCCCCTAT CGTACGTACAGCATCGCTTGCACTGAGAGCGCGGCCTCT CAGTGCAAGCGATGCTGTACGTACGATAGGGGAGGGTG GGGAGGGTGGGGAAGGTATCATGCTATGTCAATCGTGA CAGTTGCTTTTTTTTACCGGCGCTATTAGCTTCCATACCAGC TGGCAACATCCATCATGATCCGCTACTCCCA-3' |
| **7 bp Blocking oligo** | GCCGCGC |
| **RNA oligo** | GCA-UGA-UAC-CUU-CCC-CAC-CCU-CCC-CAC-CCU-CCC-C |
| **LNA oligo** | GCA-TGA-TAC-CTT-CCC-CAC-CCT-CCC-C56-668-666-6<br>For: 5 =lnA; 6 = lnC; 8 = lnT |

**Supplementary Table S1.** Sequence of the hairpins used in this assay. The colored sequences correspond to the following assemblies: blue for G4 motif, grey for the complementary sequence of the G4 motif, red for the loop, brown for the region complementary to OliBiotin, green for the region complementary to OliDBCO, yellow for the single-stranded region allowing the loading of Pif1. Oli7 is the 7-base oligonucleotide complementary to the loop.

| Name | Sequence |
|---|---|
| OliDBCO | ATT CGA AGA GCA CCA GAA AGA CCA AAA GAC ACG GTG AAG GAT TAG ACAGAA GAA GAC 3'DBCO |
| OliBiotin | 5' DualBiotin TGG GAG TAG CGG ATC ATG ATG GAT GTT GCC AGC TGG TAT GGAAGC TAA TAG CGC CGG T 3' |
| OliLoop | gcttGCACTGAGAgcgcggccTCTCAGTGC |
| Oligo5LeadG4 | GTC TTC TTC TGT CTA ATC CTT CAC CGT GTC TTT TGG TCT TTC TGG TGC TCTTCG AAT TTT TTT TGC AAC TGT CAC GAT TGA CAT AGC ATG ATA AGG GGAGGG TGG GGA GGG TGG GGA AGG ATC GTA CGT ACA GCA TC |
| Oligo3LeadG4 | aag cGA TGC TGT ACG TAC GAT CCT TCC CCA CCC TCC CCA CCC TCC CCT TATCAT GCT ATG TCA ATC GTG ACA GTT GCT TTT TTT ACC GGC GCT ATT AGC TTCCAT ACC AGC TGG CAA CAT CCA TCA TGA TCC GCT ACT CCC A |
| Oligo5LagG4 | GTC TTC TTC TGT CTA ATC CTT CAC CGT GTC TTT TGG TCT TTC TGG TGC TCTTCG AAT TTT TTT TGC AAC TGT CAC GAT TGA CAT AGC ATG ATA CCT TCC CCACCC TCC CCA CCC TCC CCT ATC GTA CGT ACA GCA TC |
| Oligo3LagG4 | aag cGA TGC TGT ACG TAC GAT AGG GGA GGG TGG GGA GGG TGG GGA AGGTAT CAT GCT ATG TCA ATC GTG ACA GTT GCT TTT TTT ACC GGC GCT ATT AGCTTC CAT ACC AGC TGG CAA CAT CCA TCA TGA TCC GCT ACT CCC A |

**Supplementary Table S2.** Sequences of oligonucleotides used to construct the hairpins of our assays

### SUPPLEMENTARY FIGURES

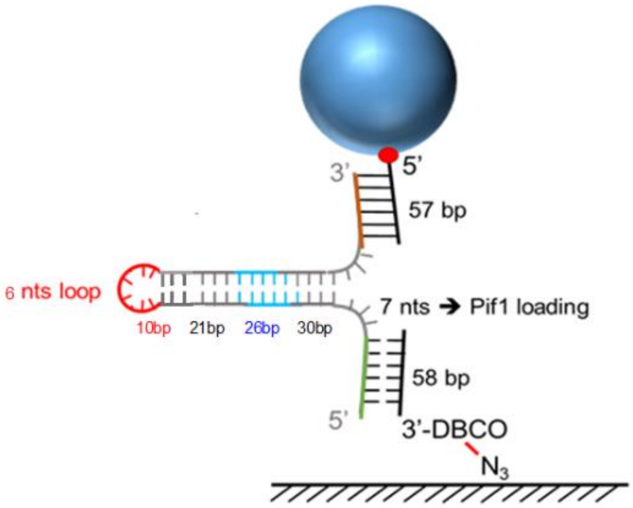


**Supplementary Figure S1. Hairpin Construct.** Hairpin construct of 87 base pairs that incorporates the 26 bp c-Myc G4 oncogene promoter sequence (c-Myc Pu27) in the middle of the hairpin. Two assays were designed, one containing the sequence in the strand before the loop (LagG4), and another after the loop (LeadG4). There are 7 nucleotides available for Pif1 to attach to the hairpin. The 5'-end of the hairpin is complementary to a 58-base 3'-DBCO modified oligonucleotide (OliDBCO), and the 3'end to a 57-base oligonucleotide (OliBiotin).

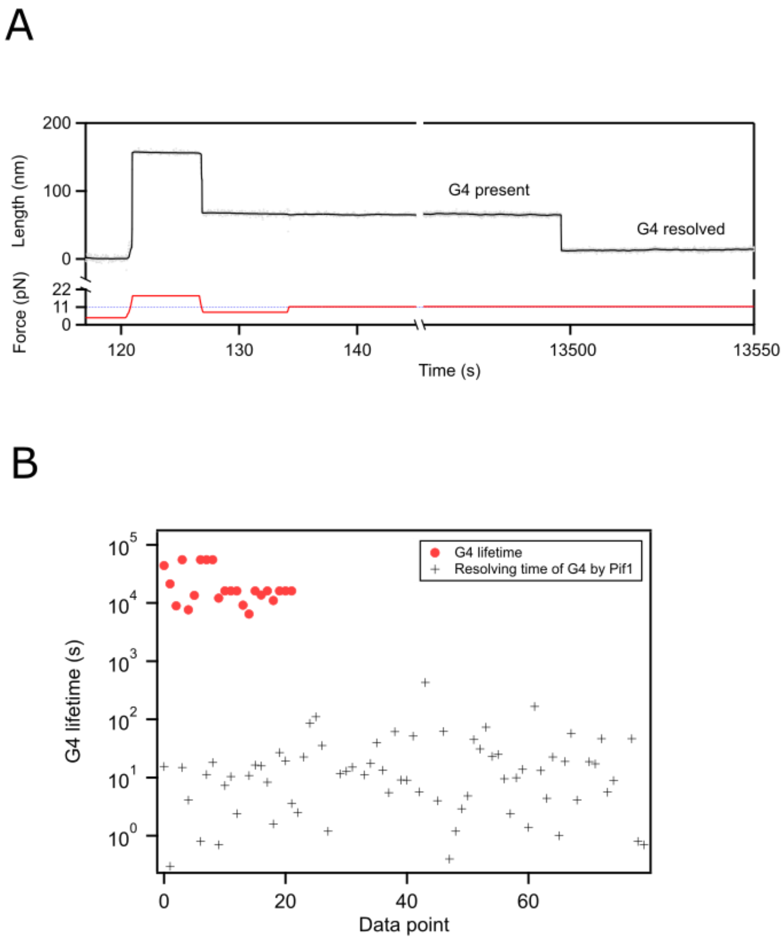


**Supplementary Figure S2. G4 lifetime. A)** The closure of the hairpin, when the force is reduced from 19 to 7 pN, is impaired by the formation of the G4 structure. This is observed as a pause. When the force is increased and maintained at 11 pN, the blockage is continuous until the G4 structure unfold, causing the fully closure of the hairpin (at 13500 s). **B)** Comparison of the lifetime of the G4 under the effects of the helicase (in log scale). If the helicase does not intervene, the G4 has an average lifetime of about **27400 ± 6300 s** . Whilst, if Pif1 collides with the G4 structure embedded in the hairpin, the structure is resolved within tens of seconds.

A

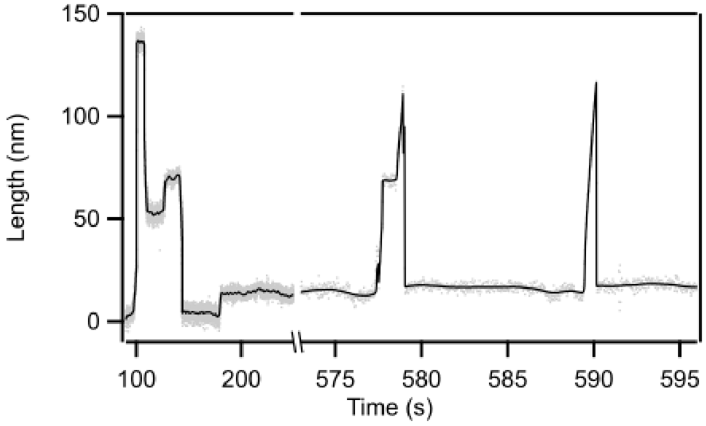


B

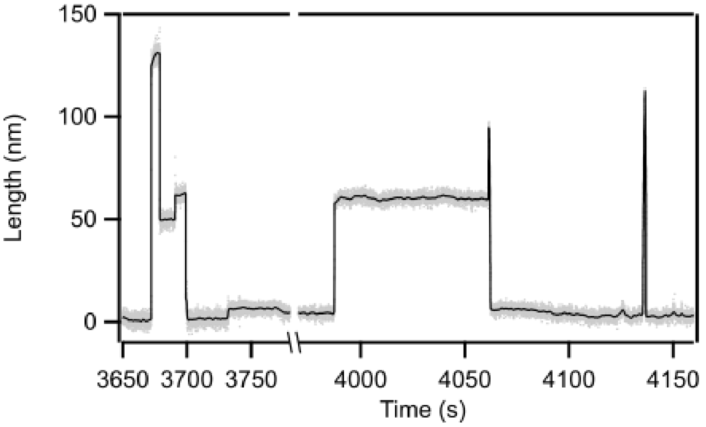


**Supplementary Figure S3. Pif1 dynamics on a LagG4 assay. A) and B)** show a typical trace where G4 has formed, as seen by a blockage at 7 pN (first cycles). When the force is kept constant at 11 pN and Pif1 binds to the hairpin, it translocates through the lagging strand until it gets stalled by the G4 structure, as seen by a blockage in the extension. The helicase resolves the structure within tens of seconds, resuming translocation. Later, another helicase binds and translocates through the whole hairpin without being blocked by the G4, indicating that it was resolved in the previous event.

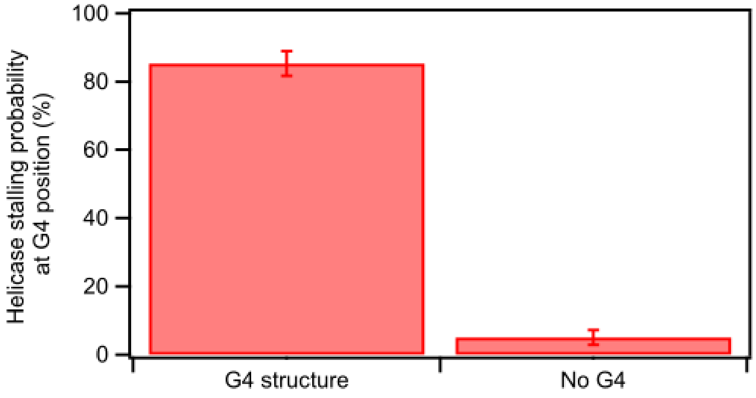


**Supplementary Figure S4. Probability of Pif1 being stalled by G4 and resolve it within the LeadG4 assay. Helicase stalling probability.** If G4 was present (G4 structure bar) within the cycles observed by a blockage at 7 pN, Pif1 gets blocked at the G4 position while translocating onto the leading strand 85% of the time. However, if G4 was not present (no G4 bar), a blockage is only observed in 5% of the traces, probably this percentage arises from mismatches in the sequence.

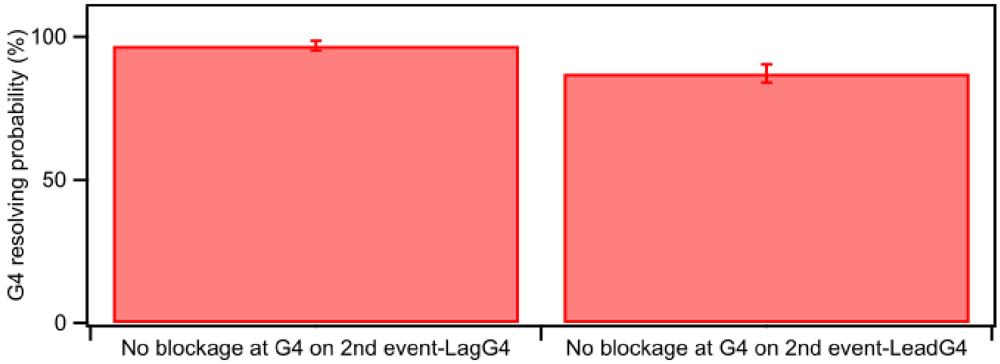


**Supplementary Figure S5. Probability of resolving G4 for LeadG4 and LagG4 assays.** IG4 resolving probability is assessed by determined the proportion of traces that do not show a blockage on the second Pif1 event. No blockage was observed in 97% and in 87% of the traces for LagG4 and LeadG4 substrates respectively.

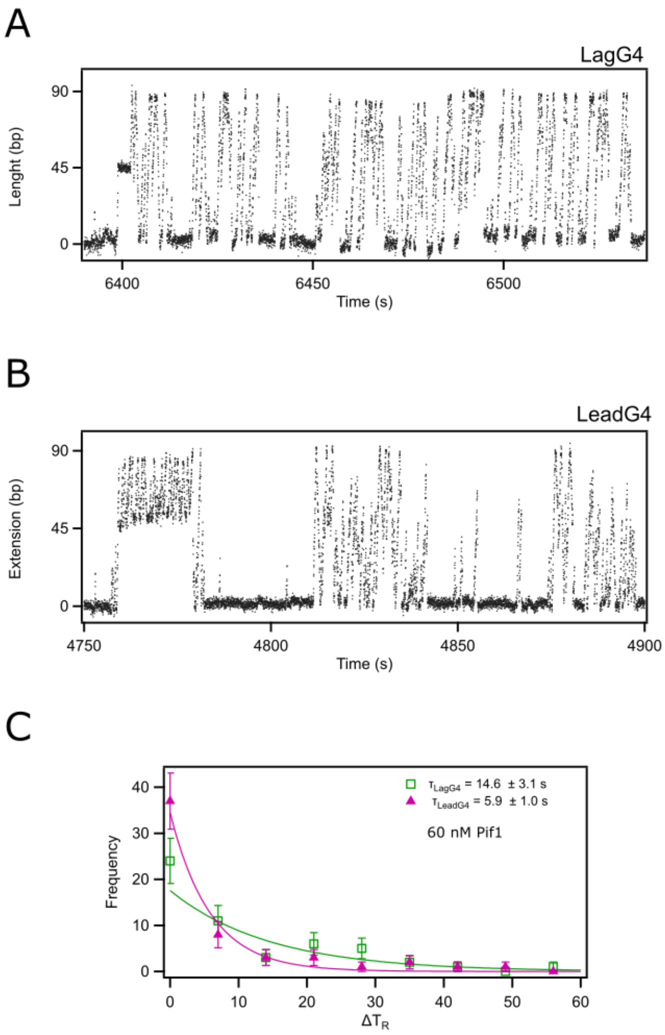


**Supplementary Figure S6. Study of the dynamics of Pif1 at a concentration of 60 nM while interacting with a DNA hairpin that has an embedded G4 structure. A) Representative trace of Pif1 when interacts with a LagG4 assay.** A characteristic blockage at the G4 position is observed on the first helicase event on the lagging strand. Translocation is resumed and the hairpin closes. Due to the high concentration the rate of a helicase binding to the hairpin increases, and the frequency of events is highly increased. **B) Representative trace of Pif1 when interacts with a LeadG4 assay.** A characteristic blockage and strand switching are observed at the G4 position, when the structure forms on the lagging strand. Once Pif1 resolves the structure and the hairpin recloses, multiple helicase events are subsequently observed within a time window of 100s. **C) Distribution of resolving time of G4 at 60nM Pif1 concentration.** resolving time shows a single exponential characterized by a time constant $\tau_{LagG4}$ (60 nM) = 14.6 ± 3.1 s and $\tau_{LeadG4}$(60 nM) =5.90 ± 1.05 s.

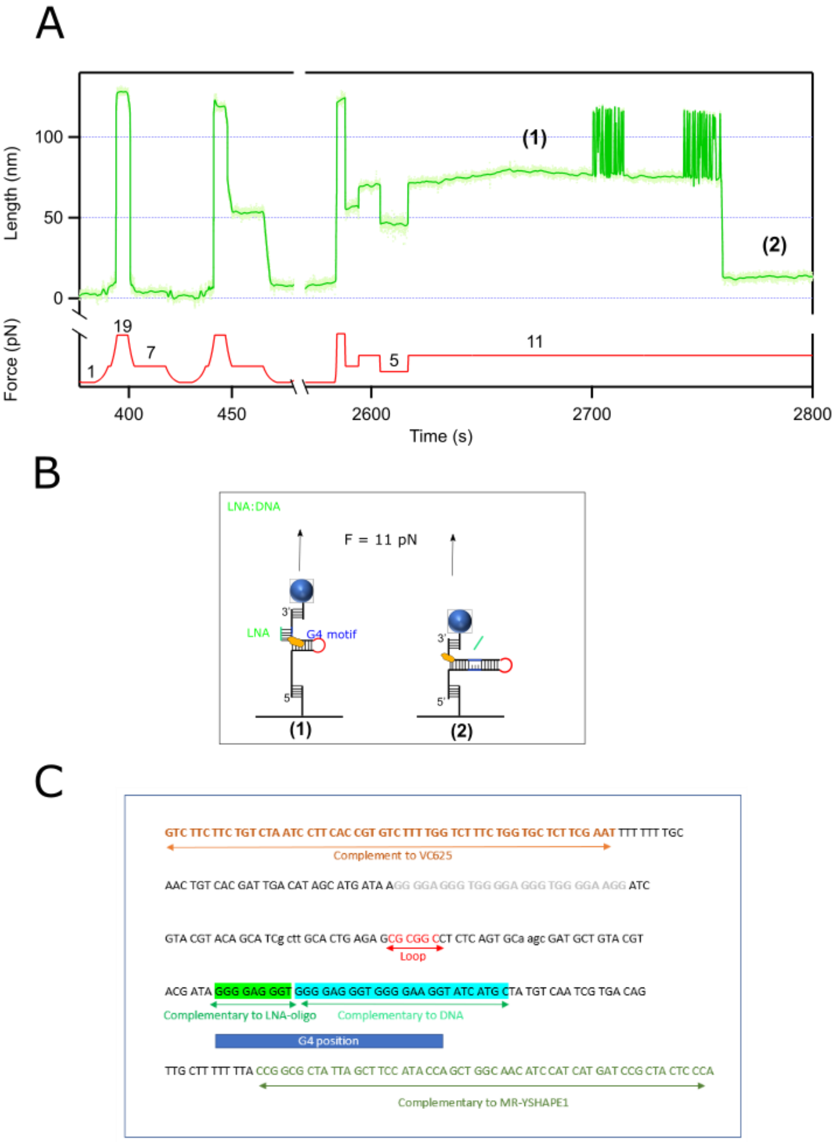


**Supplementary Figure S7. Pif1 dynamics during its interaction with a DNA:LNA hybrid complex. A) Representative trace of Pif1.** A characteristic force-extension trace showing the blockage (at 7 pN) by the LNA oligo when binds to its complementary hairpin sequence on the leading strand. At 11 pN, Pif1 is loaded into the solution. It takes several helicases to interact with the complex until the oligo is removed, as observed by the two burst of unzipping/zipping events, tens of seconds apart from each other. **B) Hairpin Sketch.** The sketches show how Pif1 interacts with the oligo within a DNA hairpin context (left), and successfully removes it (right). **C) DNA sequence.** DNA sequence highlighting the region where the LNA oligo, composed of 9 LNA and 25 DNA bases, is complementary to the hairpin, and thus will form a hybrid complex (light green). This oligonucleotide also contains the complementary sequence to the G4 motif (dark blue).

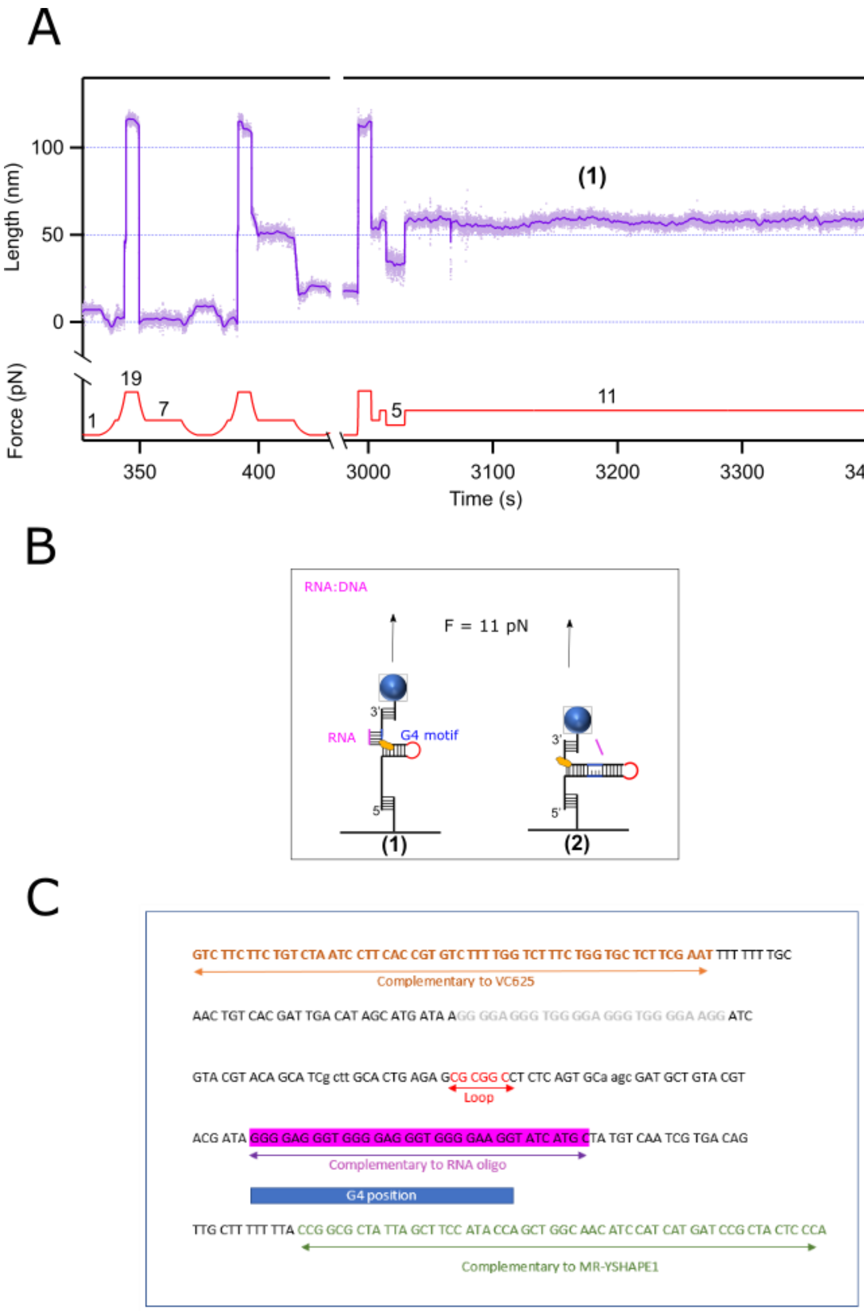


**Supplementary Figure S8. Pif1 dynamics during its interaction with a DNA:RNA hybrid complex. A) Representative trace of Pif1.** A characteristic force-extension trace showing the blockage (at 7 pN) by the RNA oligo when binds to its complementary hairpin sequence on the leading strand. At 11 pN, Pif1 is loaded into the solution, and after one translocation through the hairpin it is able to remove the RNA oligo. **B) Hairpin Sketch.** The sketches show how Pif1 interacts with the oligo within a DNA hairpin context (left), and successfully removes it (right). **C) DNA sequence.** DNA sequence highlighting the region where the RNA oligo is complementary to the hairpin and thus will form a hybrid complex (purple). Indeed, the oligo is complementary to the region of the G4 motif (dark blue).

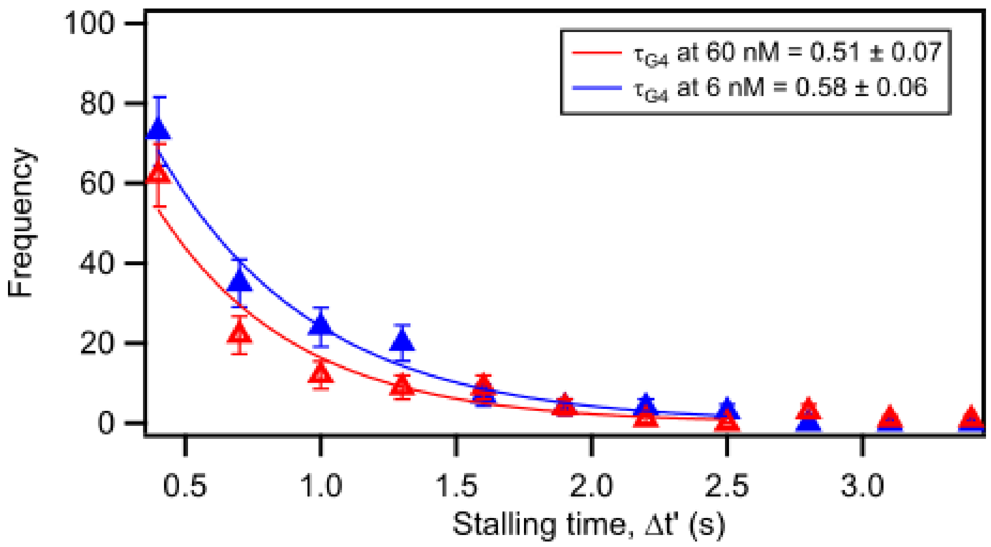


**Supplementary Figure S9. Comparison of the stalling time in the LeadG4 complex at two different Pif1 concentrations: 6 and 60 nM.** Here, the stalling time in this substrate was analyzed separately for both concentrations, to show that the differences are insignificant. Again, the data shows a single exponential characterized by a time constant $\tau_{G4}$ (6 nM) = 0.58 ± 0.06 s (blue) and $\tau_{G4}$(60 nM) =0.51 ± 0.07 s (red). Errors are sem.

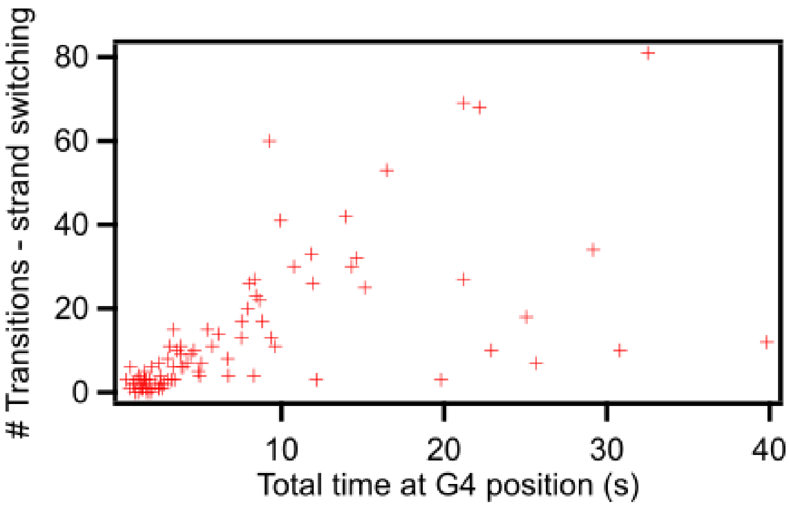


**Supplementary Figure S10.** Relationship between the total time spent at the G4 position (resolving time) and the number of strand switches. The linear relationship that the data shows infer that both phenomena are independent.

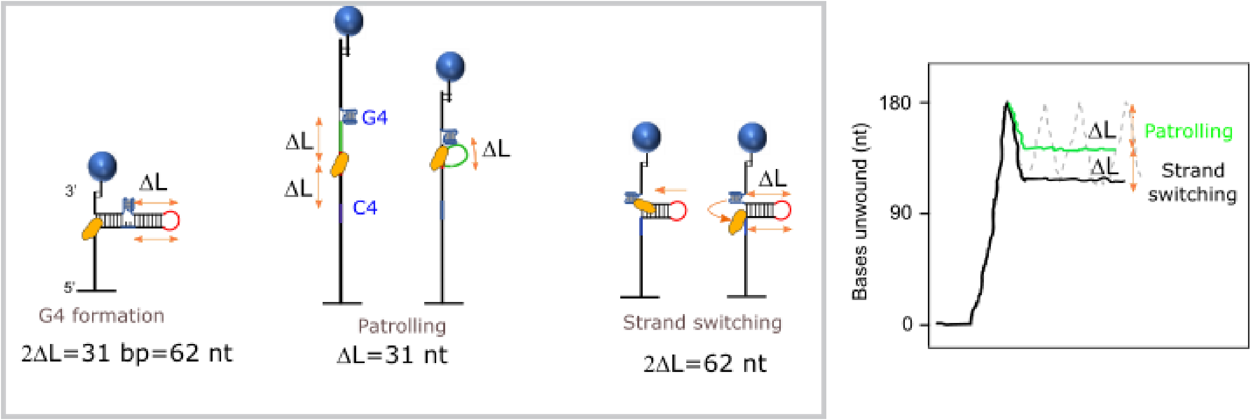


**Supplementary Figure S11. Comparison of patrolling and strand switching.** Patrolling causes the sequestering of a length dL that corresponds to the distance between the loop and the G4 structure, as seen as a shortening of the hairpin by ΔL. However, Pif1 undergoes strand switching when those 31 bp (or 62 nt), between the loop and the G4 structure, are closed so that the opposite strand is available. Thus, we measured a shortening of 2ΔL, just before Pif1 jumps onto the other strand and restarts translocation.

### SUPPLEMENTARY METHODS

Comparing translocation speeds for the different substrates.

Pif1 unwinds the double-stranded DNA along the lagging strand at a constant speed of $v_{unz1}$ = 108.37± 18.19 bp/s before interacting with the G4 structure, and resumes translocation after being stalled by the G4 at a speed of $v_{unz2}$ = 95 ± 20.13 bp/s.

We also measured both the unzipping and rezipping velocities during strand-switching transitions on the LeadingG4 substrate, and obtained very similar values within one standard deviation, of $v_{unz1}$ = 94.64 ± 9.98 bp/s and $v_{rez}$ = 113.05 ± 16.07 bp/s. Hence, these measurements imply that both translocations are the result of the same helicase process (see Table SI3).

We also analyzed the unzipping and rezipping speeds of Pif1 during strand switching in the presence of both oligonucleotides, and found for LNA a translocation speed of $v_{unz}$ = 108.22 ± 25.22 bp/s and $v_{rez}$ = 108.58 ± 20.77 bp/s, and for RNA of $v_{unz}$ = 113.39 ± 21.91 bp/s and $v_{rez}$ = 86.81 ± 32.66 bp/s for unzipping and rezipping respectively. These velocities are very similar, within one standard deviation, to the values measured on our LeadG4 assay, corroborating that the process through which Pif1 translocates along the lagging and leading strands of the hairpin in the presence of a any roadblock is the same.

Conversion procedure to transform nm into bp: shown for LeadG4 substrate:

We measured the extension at which Pif1 gets blocked by the G4 position on the leading strand, this is referred to $\Delta L$ (step 5 in Figure 2A&B). We know that the total opening of the hairpin corresponds to about 90 base pairs. Thus, we measured that quantity $\Delta L_{TOTAL}$ on our traces where Pif1 opens the whole hairpin during translocation at a constant force of 11 pN. Then, we computed a histogram and fit Gaussian function to it. Our fit showed a mean total extension of 93.0 nm with a standard deviation of 6.1 nm (Supplementary Figure S12).

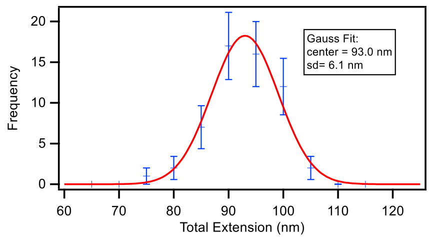

**Supplementary Figure S12. Histogram of the total extension nm units.** Histogram of the total extension of our LeadG4 hairpin at 11 pN (blue points), and the Gaussian fit (red) defining the peak at 93 nm and a spread (SD) of 6.1 nm.

We used this factor (90/93) to convert the extension of the G4 from nm to bp. And again, computed a histogram for those G4 extensions and fit it with a Gaussian (Supplementary Figure S12). This corresponds to the average position in the dsDNA hairpin at which G4 forms. In this assay, Pif1 encounters the G4 during the rezipping of the hairpin, at a stage where 49.9 ± 4.8 base pairs (data corresponds to mean and standard deviation) remain still opened.

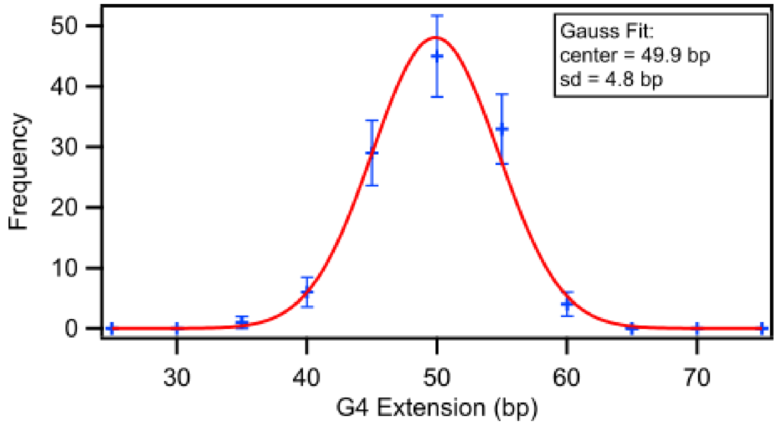


**Supplementary Figure S12. Histogram of the G4 position in base pair units.** Histogram of the converted extensions at the G4 positions (blue points), and the Gaussian fit (red) defining the peak at 49.9 bp and a spread (SD) of 4.8 bp.

At the constant force of 11 pN, Pif1 translocates through the lagging strand (unzipping) all the way to the loop and into the leading strand (rezipping). We then measured separately the unzipping ($v_{unz}$) and rezipping ($v_z$) velocities of about 122 and 101 transitions, respectively, by fitting the position of the bead with a linear function. Our results showed a mean and standard deviation of $v_{unz}$ = 97.8 ± 7.6 nm/s and $v_z$ = 116.8 ± 14.1 nm/s. We used our conversion factor determined before to provide the speed in bp/s. From the spread on the histograms of the total extension we determined an error in the conversion factor of about 6 to 8%, which probably arises from the error in the force and on the zero position. Thus, this % was added to the error on the speeds before converting them to bp/s.

This method was applied to all the speeds measured on our other substrates: LagG4, LNA:DNA, and RNA:DNA. Our values used for the conversion and the final velocities in both nm/s and bp/s are shown in Supplementary Table S3.

| **Substrate** | **Total Ext (nm)** | **Conversion bp/nm** | **Stalling position (bp)** | **Speed Unzipping $v_{unz}$ in nm/s *** | **Speed Unzipping $v_{unz}$ in bp/s** | **Speed zipping $v_z$ in nm/s** | **Speed zip $v_z$ in bp/s** |
|---|---|---|---|---|---|---|---|
| **LagG4** | 98.06 ± 8.2 | 90/98.1 | 46.3 ± 3.3 | 118.12 ± 17.43// 103.55 ± 20.32 | 108.37 ± 18.19 //95 ± 20.13 | NA | NA |
| **LeadG4** | 93.01 ± 6.11 | 90/93.01 | 49.9 ± 4.8 | 97.81 ± 7.63 | 94.64± 9.98 | 116.83±14.12 | 113.05±16.07 |
| **LNA:DNA** | 98.71 ± 7.30 | 90/98.71 | 54.3 ± 5.6 | 118.69 ± 23.64 | 108.22±25.22 | 119.09 ± 18.81 | 108.58±20.77 |

| | | | | | | |
|---|---|---|---|---|---|---|
| **RNA:DNA** | 94.49± 8.05 | 90/94.49 | 50.7± 3.9 | 119.05 ± 19.42 | 113.39± 21.91 | 91.14 ± 31.72 | 86.81± 32.66 |

**Supplementary Table S3. Translocation speeds.** Measurements on the total extension of the molecule (in nm) when the helicase unwinds the full hairpin (i.e. 90 base pairs). This extension was used to obtain a conversion factor to transform all Pif1 translocation speeds from nm/s to bp/s, for all of our substrates. * In the LagG4 substrate we obtained two unzipping speeds, one before Pif1 interacts with G4, and the other for after the stalling, when it resumes translocation.
